# Evolutionary selection on synonymous codons in RNA G-quadruplex structural region

**DOI:** 10.1101/2021.01.26.428349

**Authors:** Yuming Xu, Ting Qi, Zuhong Lu, Tong Zhou, Wanjun Gu

## Abstract

In addition to the amino acid sequence information, synonymous codons can encode multiple regulatory and structural signals in protein coding region. In this study, we investigated how synonymous codons have been adapted to the formation of RNA G-quadruplex (rG4) structure. We found a universal selective pressure acting on synonymous codons to facilitate rG4 formation in five eukaryotic organisms. While *G*-rich codons are preferred in rG4 structural region, *C*-rich codons are selectively unpreferred for rG4 structures. Gene’s codon usage bias, nucleotide composition and evolutionary rate can account for the selective variations on synonymous codons among rG4 structures within a species. Moreover, rG4 structures in translational initiation region showed significantly higher selective pressures than those in translational elongation region. These results bring us another dimension of evolutionary selection on synonymous codons for proper RNA structure and function.

## INTRODUCTION

mRNAs of protein coding genes in eukaryotic organisms not only carry the genetic information to encode amino acid sequences, but also contain multiple regulatory and structural signals ^1^. The pivot provision enabling mRNA to hold these regulatory functions is the redundancy of the genetic code that allows for many “silent” mutations at synonymous codon sites ^2^. Although synonymous nucleotide mutations do not change amino acid sequences of the encoded proteins, they can confer dramatic differences to the structure and functions of mRNA itself ^1–3^. A wealth of evidence has shown that synonymous codons are selected for optimized RNA stability ^4, 5^, proper nucleosome positioning ^6^, efficient mRNA splicing ^7, 8^, correct microRNA targeting ^9^, efficient translation initiation ^10–12^ and elongation ^13^, and proper protein co-translational folding ^14, 15^.

The RNA G-quadruplex (rG4) is a non-canonical super-secondary structure around the *G*-rich sites in RNA sequences, which consists of the stacking of G-quartets formed by the G-G Hoogsteen hydrogen bonding ^16^. Several experimental studies have shown that rG4 is ubiquitous in eukaryotes, both in untranslated regions (UTRs) and protein coding sequences (CDSs) ^17^. RG4 structures have been shown to perform many diverse and vital functions in a wide range of biological processes, such as pre-mRNA splicing ^18^, alternative polyadenylation ^19^, mRNA localization ^20, 21^, microRNA targeting ^22^, and translational regulation ^23–28^. Due to the presence of 2α-hydroxyl property, rG4 structure is more stable than DNA G-quadruplex ^29^. In a recent study, Arachchilage *et al.* has demonstrated that the most stable G4s appeared to be significantly under-represented within the CDS by the use of specific synonymous codon combinations ^30^. But, several key problems regarding synonymous codon usage and rG4 formation and evolution remain unaddressed.

Here, we looked into the evolutionary choices of synonymous codons around putative rG4 structures (pG4) at the whole transcriptome scale in multiple eukaryotic species. We asked whether there are selective pressures acting on synonymous sites at or around pG4 structures to facilitate the formation of this non-canonical super-secondary structure. If so, how synonymous codons are selected to facilitate rG4 formation? And, what are the factors that may affect this selective pressure? These analyses may help us understand the evolutionary selections acting on synonymous codons in protein coding region.

## RESULTS

### RG4 structures are selected in protein coding sequences across five eukaryotic species

We calculated the rG4 forming propensity score, *G*4*Hscore* ^31^, along the mRNA sequences using a sliding window scheme. For each pG4 structure in the transcriptome, a 30 nucleotides (nt) window was moved both upstream and downstream from the start position of the pG4 structure with a step of 30 nt, and the *G*4*Hscore* of the sequence in each window was calculated for a total of 13 windows. To estimate the background distribution of the formation propensity of rG4 structures, we randomized the mRNA sequences by shuffling synonymous codons for 1,000 times, and calculated the *G*4*Hscore* in the corresponding sliding windows. We compared the *G*4*Hscore* of the real mRNA sequence in a sliding window with that of 1,000 corresponding sliding windows in the shuffled sequences, and calculated *Z*_*G*4*Hscore*_ to assess the deviation of rG4 formation in the observed mRNA sequence from random expectation (see *Materials and Methods* for details). A positive *Z*_*G*4*Hscore*_ value means synonymous codons are selected to facilitate the formation of rG4 structures, while a negative *Z*_*G*4*Hscore*_ value means a selective pressure that prevents the formation of rG4 structures.

We performed the sliding window analysis in five eukaryotic species, including *Homo sapiens (H. sapiens)*, *Mus musculus (M. musculus)*, *Gallus gallus (G. gallus)*, *Danio rerio (D. rerio)* and *Drosophila melanogaster (D. melanogaster*). We observed a significantly positive *Z*_*G*4*Hscore*_ value in the window of pG4 structures in all five species (Figure 1). This means that synonymous codons are universally selected for the formation of rG4 structures in these five organisms. When sliding windows move to the upstream or downstream of the pG4 structures, the *Z*_*G*4*Hscore*_ values drop quickly in the flank region of pG4 structures, and oscillate around zero for sliding windows far away from the pG4 structures (Figure 1). This suggests that no selective pressures are acting on synonymous codons to facilitate or prevent the formation of rG4 structures when they are located out of the pG4 structures in the genome. When *in vitro* experimentally identified rG4 structures in human *HEK293T* cells and mouse *mESC* cells were used in the analysis, we found a similar pattern of *Z*_*G*4*Hscore*_ changes along the sliding windows (Supplementary Figure S1).

**Figure 1.**
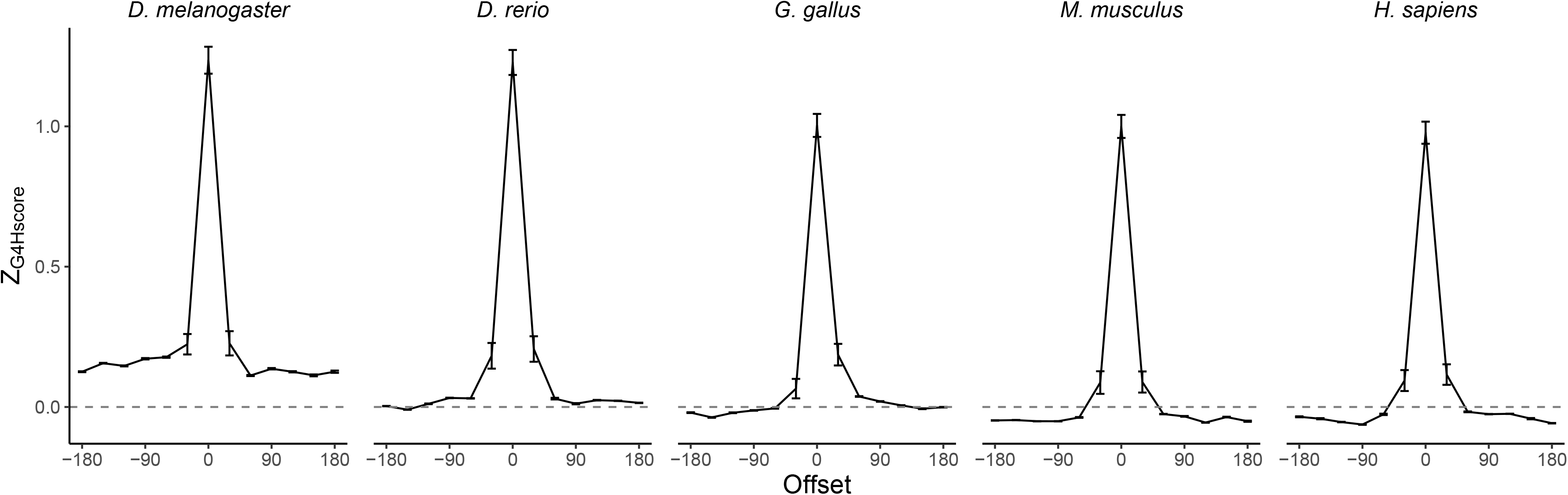
Standard error and mean value of *Z*_*G*4*Hscore*_ of 13 sliding windows in protein coding sequences in five eukaryotic species. The sliding windows are centered at pG4 structures in protein coding sequences, and moved both upwards and downwards for six windows with an offset at 30nts in length.

### G-rich or C-poor codons are selected for rG4 formation in protein coding region

To investigate how synonymous codons are selected for rG4 formation, we calculated *Z_G_* and *Z_C_* of a window of 30 nt in length for each pG4 structure in protein coding region. When comparing to *Z*_*G*4*Hscore*_ value of the same pG4 structure, we found a significant positive correlation between *Z_G_* and*Z*_*G*4*Hscore*_for pG4 structures in all five species (Figure 2). In comparison, a weaker but significant negative correlation between *Z_C_* and *Z*_*G*4*Hscore*_ of pG4 structures was also observed in all five species (Figure 2). These results suggest that synonymous codons with more *G*s and/or less *C*s are generally selected to facilitate the formation of rG4 structures in protein coding region, while the biased usage of *G*-rich codons is more relevant to rG4 formation than that of *C*-poor codons.

**Figure 2.**
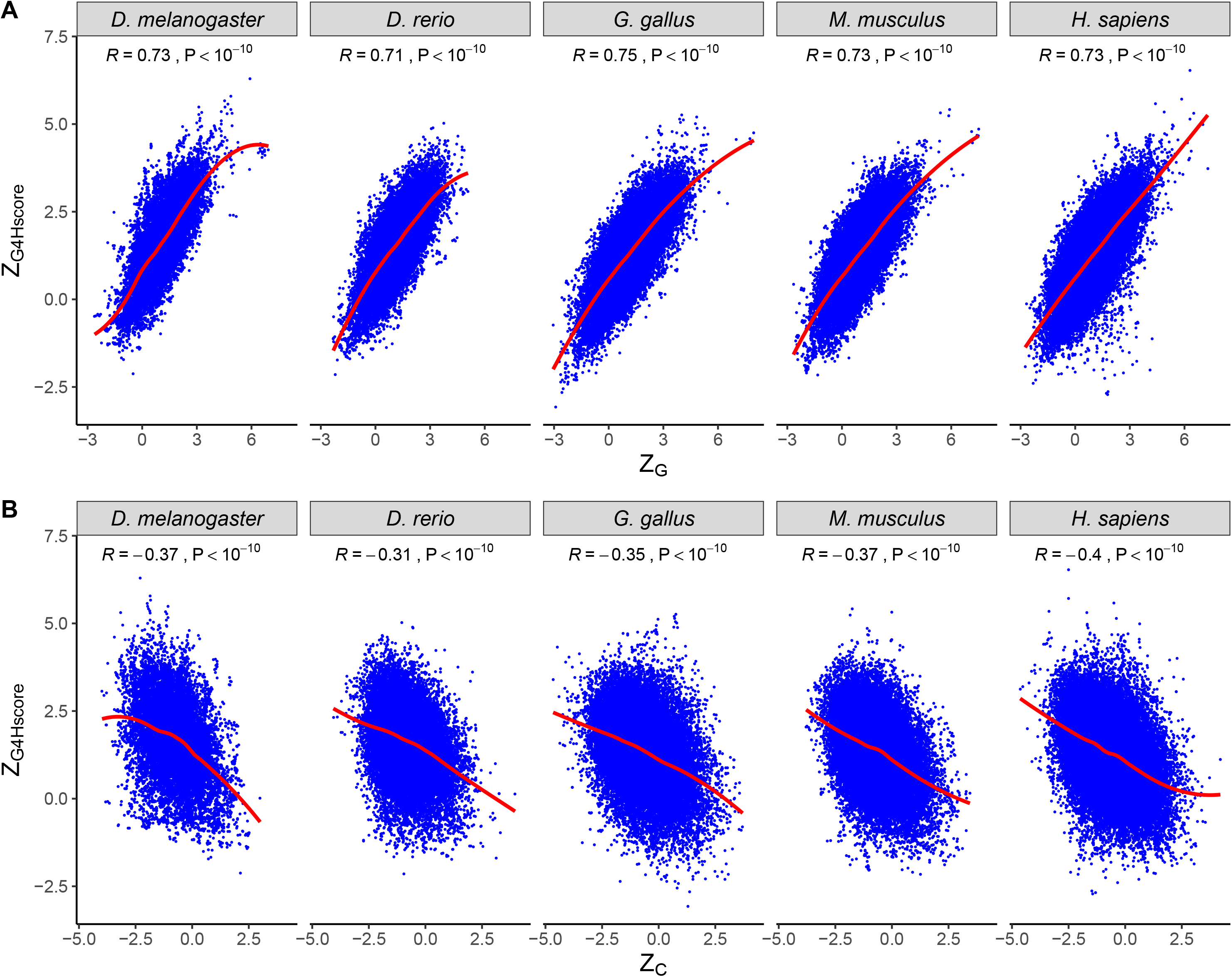
The correlations between *Z*_*G*4*Hscore*_ value and *Z_G_* **(A)** or *Z_C_* **(B)** value for each pG4 structure in protein coding region in all five eukaryotic species.

### Several features of rG4 structure’s host gene are associated with rG4 selection in protein coding region

Although the mean value of *Z*_*G*4*Hscore*_ is significantly larger than zero for all exonic pG4 structures in all five species (Figure 1), there are substantial variations among different pG4 structures within a single organism (Figures 1 and 2). To explore the factors that may affect the selective pressures on synonymous codons for rG4 formation, we considered several putative gene-level factors, including codon usage bias, evolutionary rate and nucleotide compositions of the host gene where the pG4 structure is located. We found *Z*_*G*4*Hscore*_ values of pG4 highest 5% effective number of codons (*ENC*) are significantly higher than those in genes with the lowest 5% *ENC* values (Figure 3A). This suggests that pG4 structures in genes with smaller codon usage bias are under stronger selective pressure. For the top 5% genes that have the highest *dN*/*dS* ratio, *Z*_*G*4*Hscore*_ values of pG4 structures in these genes are significantly larger than those of pG4 structures in the bottom 5% genes with the lowest *dN*/*dS* ratio (Figure 3B). When pG4 structures in genes with the highest 5% *G* content are compared to those in genes with the lowest 5% *G* content, we found pG4 structures that located in genes with the highest 5% *G* content had significantly smaller *Z*_*G*4*Hscore*_ values (Figure 3C). Moreover, *Z*_*G*4*Hscore*_ values of pG4 structures in genes with the top 5% *C* content is also statistically smaller than that of pG4 structures in genes with the bottom 5% *C* content (Supplementary Figure S2A). These results suggested that the selective pressures acting on synonymous codons are stronger for pG4 structures in genes with lower *G* nucleotides, less biased usage of synonymous codons and smaller evolutionary rate, and these differences are consistent for pG4 structures in human and mouse (Figure 3).

**Figure 3.**
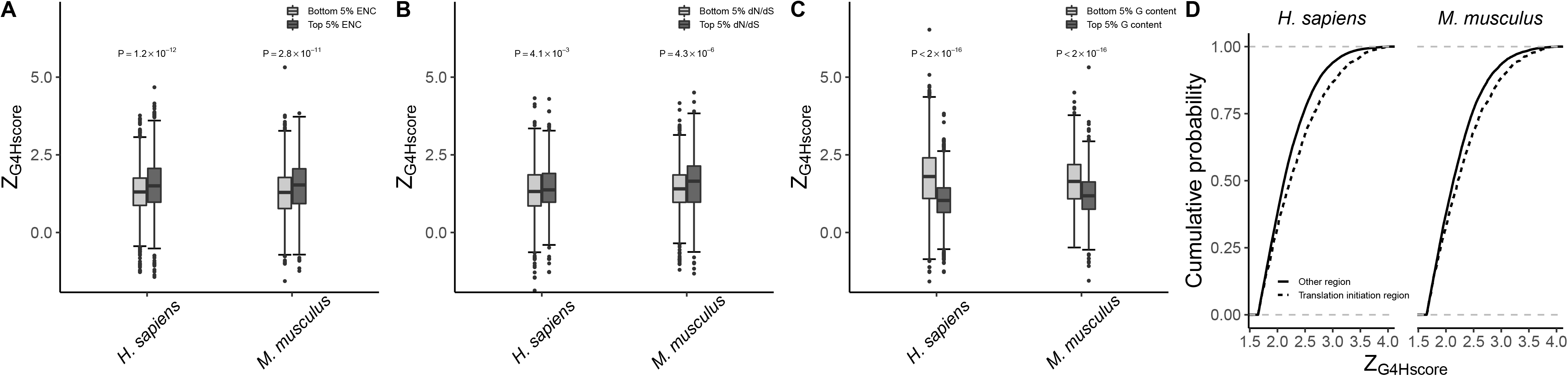
Factors that may explain *Z*_*G*4*Hscore*_ variations among rG4 structures in protein coding region of human and mouse, including *ENC* value **(A)**, *dN*/*dS* value **(B)** and *G* content **(C)** of the host gene of rG4 structure. Moreover, rG4 structures in translation initiation region have significantly higher *Z*_*G*4*Hscore*_ values than those in translation elongation region **(D)**.

### Selective pressure is different for rG4 structures in different gene regions

Other than their host genes, we also investigated *Z*_*G*4*Hscore*_ differences of rG4 structures in different gene regions. First, we grouped the pG4 structures into two categories, pG4 structures in translation initiation region (within 70nt downstream of the start codon) and those out of the translation initiation region. We found pG4 structures in the translation initiation region had significantly higher *Z*_*G*4*Hscore*_ values than those in the translation elongation region (Figure 3D). Next, we compared *Z*_*G*4*Hscore*_ values of pG4 structures near exonic splicing sites (within 60 nts of splicing sites) with those far away from the splicing sites. Although *Z*_*G*4*Hscore*_ values for pG4 structures near the splicing sites tend to be smaller than those far away from the splicing sites (Supplementary Figure S2B), the differences are not statistically significant for rG4 structures in both downstream and upstream flank regions of splicing sites. Finally, we parsed pG4 structures in microRNA (miRNA) target sites and compared their *Z*_*G*4*Hscore*_ values with those out of miRNA target region. No significant differences were observed between *Z*_*G*4*Hscore*_ values of pG4 structures in miRNA target region and those out of miRNA target sites (Supplementary Figure S2C). These results suggested synonymous codons in some functional regions, such as translation initiation region, were under different selective pressures for rG4 formation. However, rG4 structures in some regions, including miRNA target region and splicing sites flank region, did not show obviously different evolutionary selections on synonymous codons.

## DISCUSSION

In this study, we exploited the usage of synonymous codons in rG4 structural regions and saw obvious selection for the formation of rG4 structures in CDS across all five species (Figure 1). Moreover, *G*-rich and *C*-poor codons are selected in rG4 regions to facilitate the formation of rG4 structures (Figure 2). This is reasonable given the fact that sequences with more *G*s and less *C*s are more prone to form G4 structures, since *C*s may pair with *G*s to competitively form stem-loop structures instead of G4 structures ^31^. In a recent study, Archilage *et al.* ^30^ used rG4 as the model secondary structure to illustrate the impact of codon bias on secondary structure elements within the CDS of mRNAs. They observed that the most stable rG4 structures appeared to be significantly under-represented within the CDS by the use of specific synonymous codon combinations. Their findings strongly lend support to the hypothesis that rG4 structures are selectively avoided in CDS. In comparison to their conclusion, our results suggested synonymous codons, such as G-rich codons, are selected to facilitate rG4 formation when rG4 structures are located in protein coding region, although these ultra-stable RNA structures may be selectively depleted in CDS as they suggested.

We also found several features of rG4 structure’s host gene can explain the substantial variations of synonymous selection among rG4 structures within a species (Figure 3). First, genes with higher codon usage bias should have smaller chance to choose specific synonymous codons at specific rG4 structure regions, since synonymous codon usage is more biased and the repertoire of synonymous codons are smaller in these genes. Hence, we observed a reduced selection on synonymous codons in rG4 structures in genes with higher codon usage bias (Figure 3A). Second, genes with higher *dN*/*dS* ratio are more likely to experience positive selection. In comparison to rG4 structures located in conserved genes, rG4 structures in genes with higher evolutionary rate should have higher chance to be positively gained as the evolutionary outcome of recent selection. As expected, we observed a stronger evolutionary selection on synonymous codons for rG4 formation when rG4 structures are located in genes with higher evolutionary rate (Figure 3B). Third, rG4 structures located in genes with higher *G* composition are more likely to use synonymous codons with *G*s inside. Therefore, the selective pressure acting on synonymous codons in these rG4 structures may be smaller given their higher background potential to form rG4 structures (Figure 3C). However, rG4 structures in *C*-rich genes also have weaker evolutionary selection on synonymous codons (Supplementary Figure 2A). This is unexpected since *C*-rich codons are unpreferred in forming rG4 structures (Figure 2). The reason why we observed this difference is unknown, which may be partly explained by the fact that *C* content tends to be equal to *G* content at the genomic level ^32^.

In addition to above features of their host gene, we also parsed rG4 structures in several specific gene regions to investigate if synonymous codons in these rG4 structures experience significantly different selective pressures, including translational initiation region, exonic splicing sites flank region and miRNA target region (Figure 3D and Supplementary Figure S2). Translation initiation near the start codon has been reported to be the decisive process that determines translation efficiency ^10^. Our previous study revealed that reduced secondary structures were selected immediately downstream start codon to facilitate efficient translation ^11^. Several rG4 structures were observed to be located near the start codon, which can modulate gene expression by preventing the efficient recognition of *AUG* codon and blocking translational initiation ^33^. In our results, we observed that rG4 structures near start codon were under stronger evolutionary selection than other regions along the coding sequence (Figure 3D). This hints the important functions of rG4 structures in translation initiation region in gene expression regulation. In addition, several studies have reported that not only splicing sites in introns but also those in exons were influenced by the presence of rG4 structures ^18, 34–38^. The disruption of rG4 structure was found to inhibit rG4-dependent alternative splicing and cause thousands of alternative splicing events in human cell lines ^39^. However, we did not observe a significant different selective pressure when rG4 structures are located in the vicinity of exonic splicing sites (Supplementary Figure S2B). Similarly, we did not observe different selective pressures for rG4 structures in miRNA target region (Supplementary Figure S2C), although rG4 structures could play a role in miRNA-mediated gene regulation by regulating miRNA binding ^40, 41^. The reasons why we did not observe significant different selective pressures on synonymous codons in exonic splicing site flank region or miRNA binding region are not known. One possible explanation could be that functional rG4 structures near the exonic splicing sites or miRNA target region are specific to some important regulators, rather than most rG4 structures in these regions.

In conclusion, our results suggested that synonymous codons were selected to facilitate rG4 structure formation and function in protein coding genes. This brought us another dimension of evolutionary selection that acts on synonymous codons in protein coding region.

## MATERIALS AND METHODS

### Data

We downloaded the nucleotide sequences and exonic structures for all protein coding genes in five eukaryotic species, including *H. sapiens* (GRCh38.p13), *M. musculus* (GRCg6a), *G. gallus* (GRCm38.p6), *D. rerio* (GRCz11) and *D. melanogaster* (BDGP6.28), using *Ensembl BioMarts* (release 97, accessed in July, 2019) ^42^. We only considered protein coding genes with a coding sequence more than 150 nts in length. The dN and dS values of all human and mouse orthologous genes were retrieved from *Ensembl BioMarts* as well ^43^. Furthermore, we parsed miRNA target sites in protein coding regions of human and mouse genomes from *miRDB* database ^44, 45^.

To explore the factors that affect the selection for rG4 structure formation, we considered gene codon usage bias, nucleotide composition and its evolutionary rate. We used the effective number of codons (*ENC*) as the measure of codon usage bias, and calculated *ENC* values for each gene as suggested by Wright ^46^. A lower *ENC* value indicates stronger codon bias ^46^. For each gene, we also calculated nucleotide compositions, including *G* content and *C* content. Moreover, we calculated the ratio of *dN* and *dS* values for each gene, and used the *dN*/*dS* value as the measurement of evolutionary rate. A *dN*/*dS* value close to zero means the gene is conserved between human and mouse.

### rG4 structure in protein coding region

To locate rG4 structures in the protein coding region, we used the *G4Hunter* algorithm ^31^ to systematically search for potential rG4 forming sites in protein coding sequences in all five species. We ran *G4HunterApps* with a window size at 25 nt and a cutoff at 1.2 ^47^ (https://github.com/LacroixLaurent/G4HunterMultiFastaSeeker) to identify all potential rG4 structures (pG4) in all protein coding sequences. We chose *G4Hunter* algorithm with these two parameters, since a comprehensive evaluation of computational methods for rG4 prediction has suggested that *G4Hunter* has the best performance in predicting G4 structures ^48^. We identified 65,562, 42,153, 38,311, 25,194, and 14,184 rG4 structures in protein coding sequences for *Homo sapiens*, *Mus musculus*, *Gallus gallus*, *Danio rerio*, and *Drosophila melanogaster*, respectively. To validate the observations that we found in predicted rG4 structures, we also downloaded experimentally derived rG4 sites in human *HEK293T* cells and mouse *mESC* cells by high throughput RT-stop techniques from supplemental materials of Guo *et al.* ^49^.

### mRNA Randomization

If the choice of synonymous codons influences the formation of rG4 structures in coding sequences, the *G4Hscore* of mRNA sequences in naturally occurred pG4 region should be statistically different from that of those randomized sequences. Thus, we randomly shuffled synonymous codons in the coding sequence, keeping the amino acid sequence, *GC* composition and codon usage bias the same. For each CDS sequence, the shuffling process was repeated 1,000 times to obtain a set of randomized artificial sequences. The *G4Hscore* of each 30nt window in the native CDS sequence and each permutated sequence were calculated using *G4Hunter* algorithm ^31^. The difference of G4 forming potential between the native sequence and the randomized sequences was determined by calculating the *Z*-score of *G4Hscore* for each sliding window by

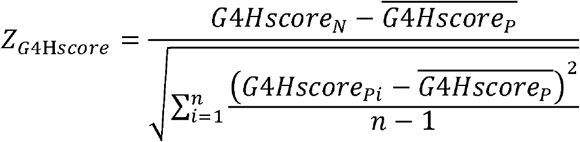

Here, *G*4*Hscore_N_* is the *G4Hscore* for the native sequence in the window, *G*4*Hscore_Pi_* is the *G4Hscore* of the corresponding window of the *i*^th^ randomized sequence, and 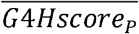 is the mean of *G*4*Hscore_Pi_* over all randomized sequences. The variable *n* represents the total number of randomized sequences, which is 1,000 here.

Similarly, the difference between the *G* (or *C*) compositions of the native sequence and the randomized sequences was evaluated. The *Z*-score of *G* (or *C*) content (*Z_G_* or *Z_C_*) for each sliding window can be calculated by

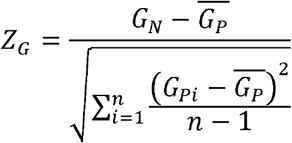

and

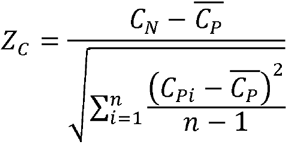

The meanings of *G_N_* (*C_N_*), *G_Pi_* (*C_Pi_*) and 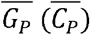 are analogous to those of *G4Hscore*, but refer to *G* or *C* content instead of G4 forming potential.

## ACKNOWLEDGEMENTS

This work was funded by grants from National Key R&D Program of China (2018YFC1314900, 2018YFC1314902), National Natural Science Foundation of China (61571109), and the Fundamental Research Funds for the Central Universities (2242017K3DN04).

## SUPPLEMENTARY FILES

**Supplementary Figure 1**. Standard error and mean of *Z*_*G*4*Hscore*_ of 13 sliding windows in protein coding sequences in human and mouse. The sliding windows are centered at *in vitro* experimentally validated rG4 structures in protein coding sequences, and moved both upwards and downwards for six windows with an offset at 30nts in length.

**Supplementary Figure 2**. **(A)** *C* content of the host gene of rG4 structure may not explain *Z*_*G*4*Hscore*_ variations among rG4 structures in protein coding region of human and mouse. In addition, when comparing to rG4 structures in other protein coding regions, rG4 structures in the flank region of exonic splicing sites **(B)** and miRNA target region **(C)** did not show significantly different *Z*_*G*4*Hscore*_ values.

