## Supplementary figures and images for "Evolutionary selection on synonymous codons in RNA G-quadruplex structural region"

### supplementary figure 1

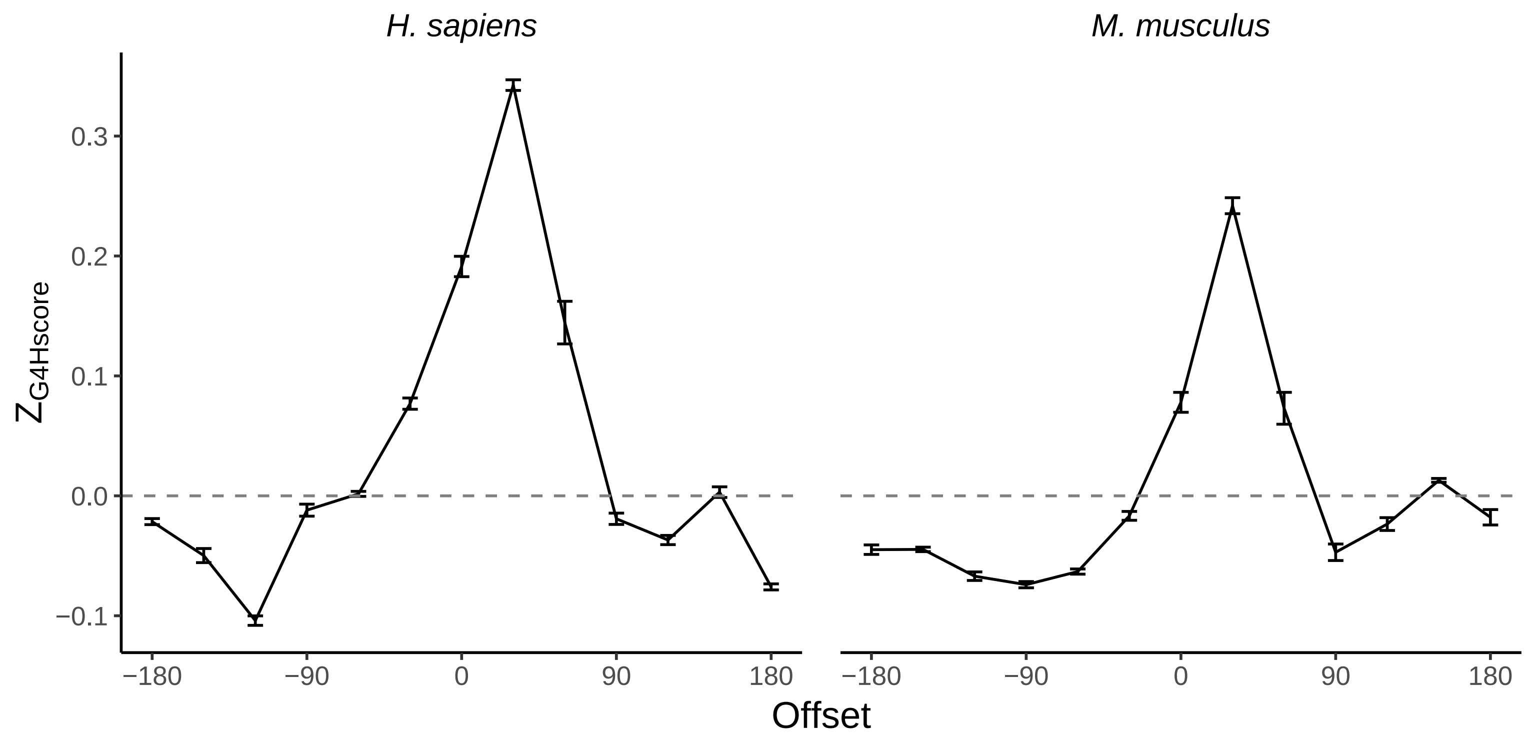

### supplementary figure 2

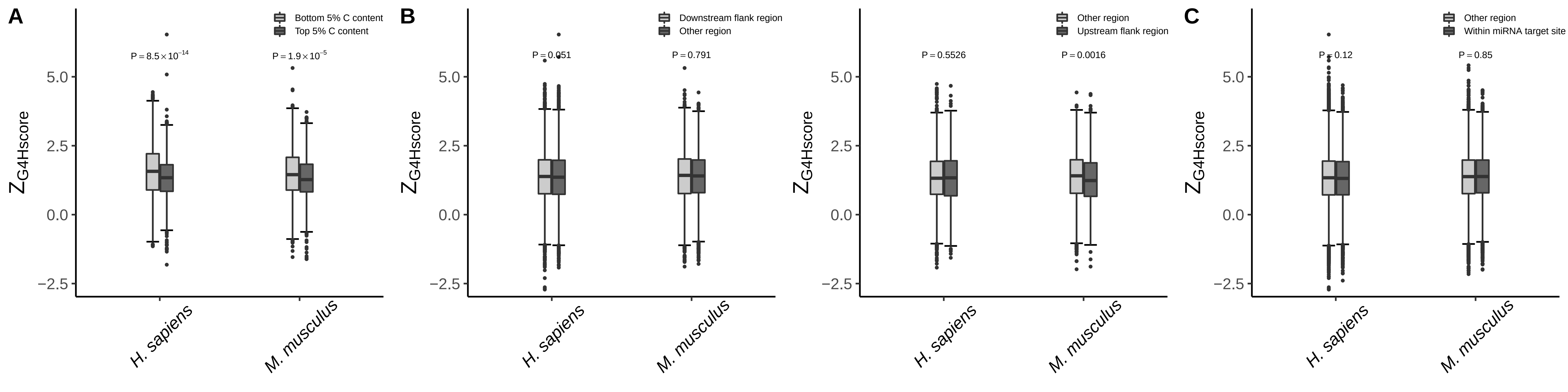
